# Understanding neural circuit development through theory and models

**DOI:** 10.1101/121574

**Authors:** Leonidas M. A. Richter, Julijana Gjorgjieva

## Abstract

How are neural circuits organized and tuned to achieve stable function and produce robust behavior? The organization process begins early in development and involves a diversity of mechanisms unique to this period. We summarize recent progress in theoretical neuroscience that has substantially contributed to our understanding of development at the single neuron, synaptic and network level. We go beyond classical models of topographic map formation, and focus on the generation of complex spatiotemporal activity patterns, their role in refinements of particular circuit features, and the emergence of functional computations. Aided by the development of novel quantitative methods for data analysis, theoretical and computational models have enabled us to test the adequacy of specific assumptions, explain experimental data and propose testable hypotheses. With the accumulation of larger data sets, theory and models will likely play an even more important role in understanding the development of neural circuits.

## Introduction

Neural systems are tuned to enable the efficient and stable processing of information across different brain regions and to generate robust behaviors. This requires a balance between flexibility, to learn from and adapt to new environments, and stability, to ensure reliable execution of behavior. Generating systems with this double property is a non-trivial challenge and requires a prolonged period of development when multiple mechanisms are coordinated in a hierarchy of levels and timescales to establish a rich repertoire of computations. Studying this process is of fundamental importance for the understanding of normal brain function and the prevention, detection and treatment of brain disorders, including intellectual disabilities, autism, bipolar disorder, schizophrenia and epilepsy.

The developing brain is not merely an immature version of the adult brain. Even before sensory experience begins to sculpt connectivity, a diversity of mechanisms and structures unique to development characterize the self-organization into functioning circuits. Technological advancements in experimental techniques have made feasible to record and manipulate a number of circuit components. In parallel, data analysis techniques and theoretical and computational models have enabled us to synthesize experimental data from multiple systems and to derive key principles for how neural circuits are built and organized into functional units, which can adapt to, learn from and discriminate different sensory stimuli.

We highlight recent theoretical work on neural circuit organization during early stages of development (late prenatal or early postnatal) before sensory organs mature. We focus on activity-dependent mechanisms governing this process, after neuronal differentiation and migration have taken place, and use the visual system and the immature (undifferentiated) cortex as examples. By describing theoretical and modeling approaches for spontaneous activity generation, developmental refinements of connectivity and intrinsic single neuron properties, and the emergence of computations, we highlight the success of theoretical models to analyze different mechanisms and propose new hypotheses of neural circuit development.

## Models of topographic map formation in the visual system

The initial stages of circuit development consist of establishing precise patterns of connectivity guided by matching molecular gradients and axonal targeting. One of the best studied models of organization of neural circuit connectivity are topographic maps in the visual system, whose orderly structure has made them an accessible model system for both theory and experiment. Retinotopic maps between the retina and higher visual centers, including the superior colliculus (SC), the lateral geniculate nucleus (LGN) and the cortex, have been the focus of intense study, elucidating general principles underlying neural circuit wiring [1–6, 7^•^, 8^••^]. Most models assume that topographic maps are formed by the interaction of molecular guidance cues, such as Ephs/*ephrins* (reviewed in [5, 9]), and are subsequently refined by spontaneous neural activity. We highlight three aspects of recent progress on map formation before we explore more computational aspects of development.

Recent models simulate not only the final map, but the **entire temporal evolution of map formation** from a combination of mechanisms: retinal axons initially arborize stochastically in the target region, synaptic connections are subsequently refined by Hebbian activity-dependent plasticity and regulated in strength through competition for a common source [10, 11, 12^••^]. Despite the success of these models in reproducing experimental results, one disadvantage is their complexity – structure emerges from many interacting mechanisms making it difficult to infer which of the resulting properties are the product of any of the model ingredients; another disadvantage is their reproducibility since they take days to simulate.

With the accumulation of experimental data from normal and mutant animals, **new quantitative analysis methods of maps** have also been developed. One example is the ‘Lattice Method,’ which enables a quantitative assessment of the topographic ordering in the one-to-one map between two structures [13]. Fitting models to data from different types of mutants has suggested that that activity-dependent and molecular forces most likely act simultaneously (rather than sequentially) and stochastically to give rise to ordered connections as observed experimentally [13, 14].

To compare and unify different models aimed at capturing specific aspects of map formation, new frameworks now support the **unbiased and quantitative testing of computational models** on available data from the mouse retinocollicular system [15^••^]. These enable us to go beyond comparing model output to known perturbations and towards predicting how these models would respond to novel manipulations. Such approaches are especially useful when several different models are equally consistent with existing data [16^••^]. Despite the success in modeling map formation, the challenge remains to integrate maps with the emergence of other, more functional aspects of development.

## Spontaneous activity: transient features and computational implications

Before the onset of sensory experience, many developing circuits can generate neural activity spontaneously. Spontaneous activity regulates many developmental processes, including neuronal migration, ion channel maturation and the establishment of precise connectivity [9, 17, 18]. In the retina, spontaneous activity is generated before the retina responds to light, and has been commonly implicated in the refinement of retinotopic maps between the retina and SC or LGN [18, 19]. The retina generates complex spatiotemporal waves of spontaneous activity during the first two weeks of postnatal development (in rodents) ([20], for models see [21, 22]). These propagate through the visual pathway to the SC, the thalamus, and the visual cortex [23, 24, 25^•^], which are themselves spontaneously active [26^•^, 27]. We review several transient cellular properties and transient structures contributing to the generation and propagation of spontaneous activity in the cortex, and subsequently examine its computational implications and role in the refinement of connectivity to downstream targets.

Developing neurons express a unique configuration of ion channels and receptors to mediate specific patterns of spontaneous activity, which may be incompatible with the information-processing functions of mature neurons [17]. In the developing mouse cortex, the proportions of the two main spike-generating conductances (sodium and potassium) of single neurons change during the first postnatal week. This biophysical change enables single neurons to gain an ability to dynamically adjust their response range to the size of stimulus fluctuations [28]. This property is termed ‘gain scaling’ and can be extracted by building linear-nonlinear (LN) models from the response of single neurons to random noisy stimuli (Fig. 1A,B)[28]. Gain scaling in more mature neurons supports a high rate of information transmission about stimulus fluctuations in the face of changing stimulus amplitude, and is absent in immature neurons (Fig. 1C) [29^••^].

**Figure 1:**
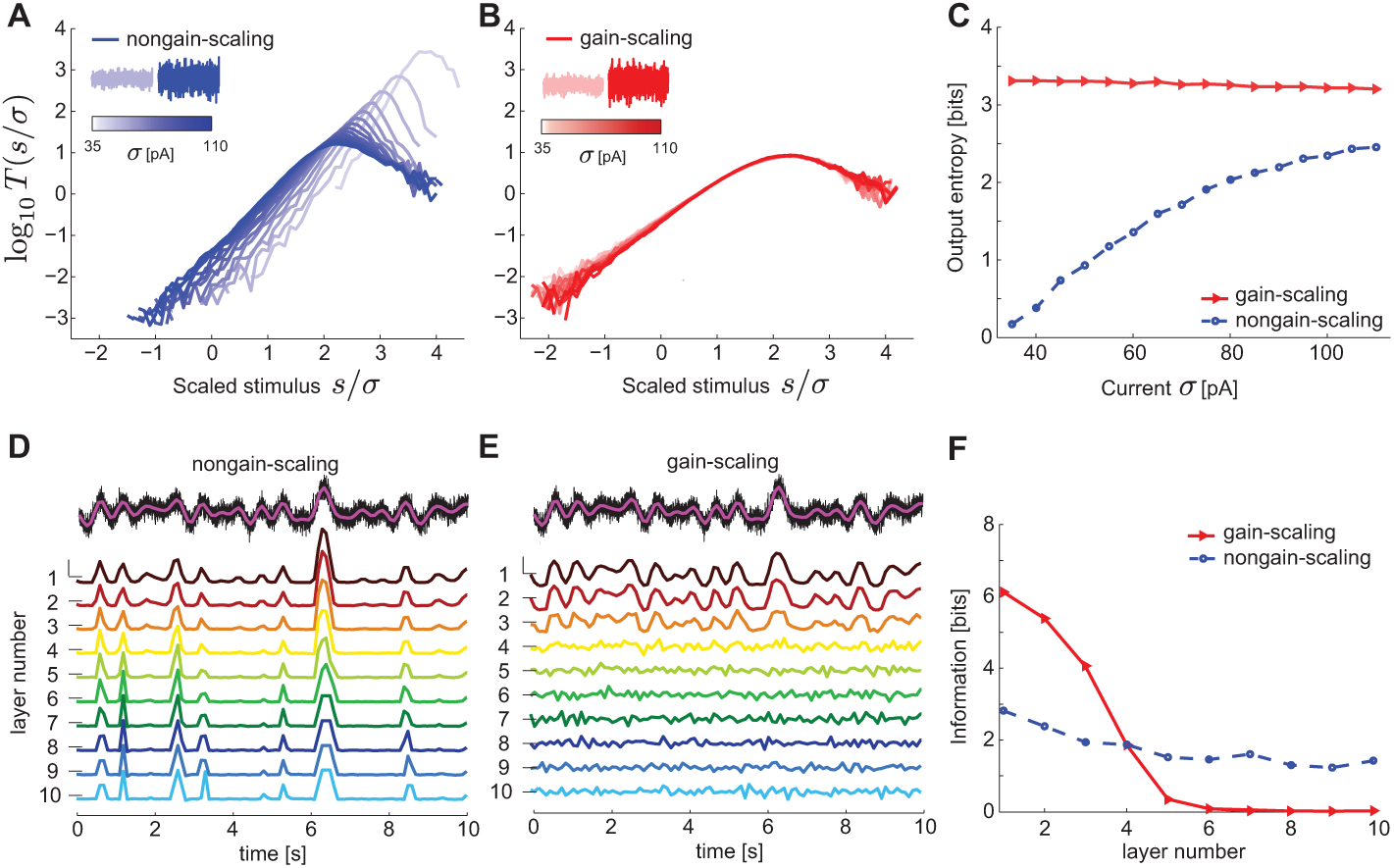
**A,B.** The nonlinearities in the LN model framework for a nongain-scaling (A) and a gain-scaling (B) Hodgkin-Huxley neuron stimulated with white noise with a range of variances *σ*^2^. The lack of gain scaling corresponds to cortical neurons recorded around birth, while the ability to gain scale matches cortical neurons after the first postnatal week. The nonlinearities were computed using Bayes’ rule: *T*(*s*) = *P*(spike|*s*)/*r* = *P*(*s*|spike)/*P*(*s*) where *r* is the neuron’s mean firing rate and s is the stimulus filtered by the spike-triggered average [29^••^]. The nonlinearities of gain-scaling neurons overlap over a wide range of *σ*. **C.** The output entropy as a function of the amplitude of fast fluctuations, *σ*, measures the information about fast fluctuations. **D,E**. Peristimulus time histograms (PSTHs) from each layer in feedforward networks of nongain-scaling (D) vs. gain-scaling (E) neurons showing the propagation of a slow-varying input (magenta, top) in the presence of background fast fluctuations (black, top). PSTHs were normalized to mean 0 (horizontal line) and variance 1 (vertical scalebar = 2). **F.** Mutual information about the slow-varying input (D,E magenta, top) transmitted by the two networks in D and E. Figure adapted from [29^••^].

These single neuron changes in gain scaling during development can generate very different dynamics at the network level [29^••^]. The lack of gain-scaling early in development (around birth) allows slow activity transients to propagate through model cortical networks (Fig. 1D). This can be related to the ability of the cortical network to propagate spontaneous waves at birth. The emergence of gain scaling a week later when spontaneous waves disappear, makes the networks better suited for the efficient representation (but not propagation) of information on fast timescales relevant for sensory stimuli (Fig. 1E) [29^••^]. These different abilities of the two networks to transmit slow stimulus fluctuations, can be captured in the mutual information between the slow stimulus and the average network response (Fig. 1F). This example demonstrates that single neuron properties can influence network dynamics, thus bridging the gap between two levels of description, and makes predictions for the information processing capabilities of these networks which can be evaluated in experimental data.

To model cortical spontaneous activity in more biologically realistic set ups requires that spontaneous transients are endogenously generated by the networks themselves, rather than provided as input to the network (as in Fig. 1E,F). To determine the source of these transients consistent with data, Baltz and colleagues proposed three different models implicating intrinsic bursts, spikes or accumulation of random synaptic input [30]. Although all models could generate propagating spontaneous events, networks where single neurons produced intrinsic bursts were most consistent with *in vitro* recordings of spontaneous network activity [30]. Barnett and colleagues distributed intrinsically bursting neurons along a gradient in a model network with recurrent synaptic connectivity and local gap junctions. The gradient of intrinsic bursting was sufficient to capture the direction of wave propagation in cortical slices [31^•^]. The models also predicted that wave activity persists near the site of initiation even after a wave has passed, which was confirmed experimentally [31^•^].

Other transient network features are also prominent in developing circuits. The depolarizing action of GABA in immature circuits (reviewed in [32, 33]) is an example of a transient developmental feature which several models have utilized for the propagation of spontaneous activity [30, 31^•^] – this feature seems to be important to support spontaneous activity in networks where immature neurons have high excitability thresholds, and weak and unreliable connectivity. The subplate is a transient structure with relatively mature properties, which serves as a scaffold to establish strong and precise connectivity between the thalamus and cortex and then disappears [34, 35]. As a third example, we mention the transient excitatory feedback connectivity between the thalamic reticular nucleus (TRN), thalamus and visual cortex, which appears necessary for the generation of feedforward connectivity along the developing visual pathway [36^••^]. The TRN and the subplate have so far not been modeled, except for a circuit subplate model with a single neuron at each relay stage (thalamus, subplate and cortex) [37], leaving open the question of how these transient structures support the organization of large neural circuits with multiple convergent and divergent connections.

Network models incorporating these transient features could shed light on how developing circuits become spontaneously active even when cellular properties are immature and connectivity is still forming. Models offer the advantage of studying the action of any mechanism independently from the rest, allowing us to identify the relative influence of each, as has been done with ion channel distributions and intrinsic excitability gradients.

## Linking neural activity to the refinement of connectivity

How can developmental activity patterns (spontaneous or sensory-evoked) guide synaptic connectivity refinements? Detailed analysis of the spatiotemporal structure of activity can provide insights into the nature of the operating rules of synaptic plasticity. During early development, patterns of spontaneous activity are ‘sluggish’ and characterized by long lasting events (bursts, spindle bursts, and calcium-dependent plateau-potentials) that have correlation timescales on the order of hundreds of milliseconds [23, 26^•^, 38–41]. Therefore, it is natural to assume that the plasticity rules that translate these patterns into circuit refinements should operate over long timescales.

Theoretical studies of phenomenological plasticity rules have helped us understand the implications of different spatiotemporal structure of activity on the temporal evolution of connectivity. The activity patterns are typically interpreted into functional synaptic changes and circuit organization through Hebbian rules that use pre- and postsynaptic activity to increase or decrease synaptic strength. One of the best studied forms of Hebbian plasticity in theoretical models is Spike-Timing-Dependent Plasticity (STDP), where potentiation and depression are induced by the precise timing and temporal order of pre- and postsynaptic spikes [42]. Because this classical STDP rule integrates input correlations on the order of tens of milliseconds – much faster than firing in development – more appropriate rules for developmental refinements have been analyzed. These include STDP rules which integrate more spikes or long temporal averages of the membrane potential (e.g. triplet STDP, voltage STDP) [43–45] and burst-based rules (e.g. BTDP) which evoke synaptic potentiation and depression based on the overlap (but not order) of bursts of spikes [38, 39]. These plasticity rules have been studied in feedforward model networks where an array of input units projects to a single postsynaptic neuron, successfully explaining the emergence of various developmental receptive field features, including eye-specific segregation [38], ON-OFF segregation [39], and direction selectivity [45, 46].

A recent study connected these mechanistic connectivity refinements from known plasticity rules to normative models for the emergence of receptive field structures [47^•^]. By developing the concept of *nonlinear Hebbian learning*, the theory simultaneously satisfied the requirements for final receptive field structure and the mechanisms for its development [47^•^]. This type of learning arises from the combination of plasticity with a neuron’s input-output function and can be implemented by sparse coding and independent component analysis [48, 49]. Coupling neurons into recurrent networks resulted in the development of diverse receptive fields through nonlinear Hebbian learning, which was necessary to represent the entire space of possible stimuli. Going one step further, it would be interesting to extend this theory to link bottom-up and top-down approaches for plasticity in recurrent synapses.

These theoretical studies drive synaptic refinements based on low-order correlations measured in spontaneous activity and early evoked responses. However, developmental activity patterns contain much more structure on several temporal and spatial scales, and activity itself refines during brain maturation [25^•^, 26^•^, 50^•^]. At the same time, these activity-dependent refinements interact nontrivially with molecular mechanisms as discussed earlier [10, 12^••^]. A future challenge is to determine how more complex activity patterns could shape network connectivity and sensory representations in models which are still analytically tractable.

## The emergence of systems-level organization

One major challenge for modeling the implications of realistic developmental patterns is plasticity in recurrent networks of spiking neurons; this problem has been recently approached in numerical simulation studies. Clopath and colleagues introduced a biologically motivated plasticity rule for spiking neurons, voltage STDP, to model plasticity in spiking recurrent networks with different input configurations [44]. Although the assumed input patterns were abstract groups of correlated neurons, unrelated to experimentally recorded activity, plasticity in the networks generated different functional network structures including synfire chains and self-connected assemblies accompanied by prevalence of bidirectional connections [44].

Voltage STDP was also applied to a developmental scenario of the emergence of functional specificity of recurrent connections among similarly-tuned neurons in mouse V1 [51^••^]. This specificity of recurrent connections only emerges after eye-opening, building on feature preference of individual neurons which is already present at eye-opening [51^••^]. To capture the additional aspect of feature preference before eye-opening, the same plasticity rule was implemented at feedforward synapses preceding any recurrent plasticity. The presence of gap junctions among specific cortical neurons was used to establish initial selectivity biases that were eventually amplified by recurrent plasticity and redistribution of recurrent synaptic connections [51^••^]. Therefore, the action of a single phenomenological plasticity rule successfully captured the experimentally observed sequence of developmental events from feedforward feature preference acquisition, to the emergence of recurrent connection specificity among similarly-tuned neurons.

Sadeh and colleagues studied a comparable process in large recurrent networks of spiking neurons with balanced excitation and inhibition, where the dominant input to a neuron is not feedforward but comes from the local recurrent network into which the neuron is embedded [52^•^]. This recurrent input sharpened the initially weak orientation selectivity of single neurons. Plasticity at both recurrent excitatory and inhibitory synapses produced adult connection specificity [52^•^]. In addition to sharpening orientation selectivity, the neurons also sparsified their responses as observed experimentally around eye opening [53, 50^•^]. One caveat of both models [51^••^, 52^•^] is that they do not explicitly represent orientation selectivity: the emergence of this feature selectivity is realized by the selective potentiation of feedforward inputs from a group of correlated neurons. Related models, however, can give rise to biphasic, oriented receptive fields localized in space under certain conditions [54^•^].

More broadly, preferentially strong connectivity among groups of neurons in recurrent network models with balanced excitation and inhibition can emerge without reference to the feature preference (or sensory tuning) of these neurons [55^•^, 56, 54^•^]. These preferentially connected groups are called *Hebbian assemblies*; the attractor dynamics they can give rise to in networks [54^•^, 57] could be the substrate of different neural computations, including predictive coding through the spontaneous retrieval of evoked response patterns (Fig. 2) [54^•^, 55^•^, 56] and decreased variability during sensory stimulation [55^•^]. Interestingly, in some of these models recurrent attractor dynamics and biphasic, oriented receptive fields localized in space emerge only when the networks are trained with natural image stimuli, but not with white noise [54^•^].

**Figure 2:**
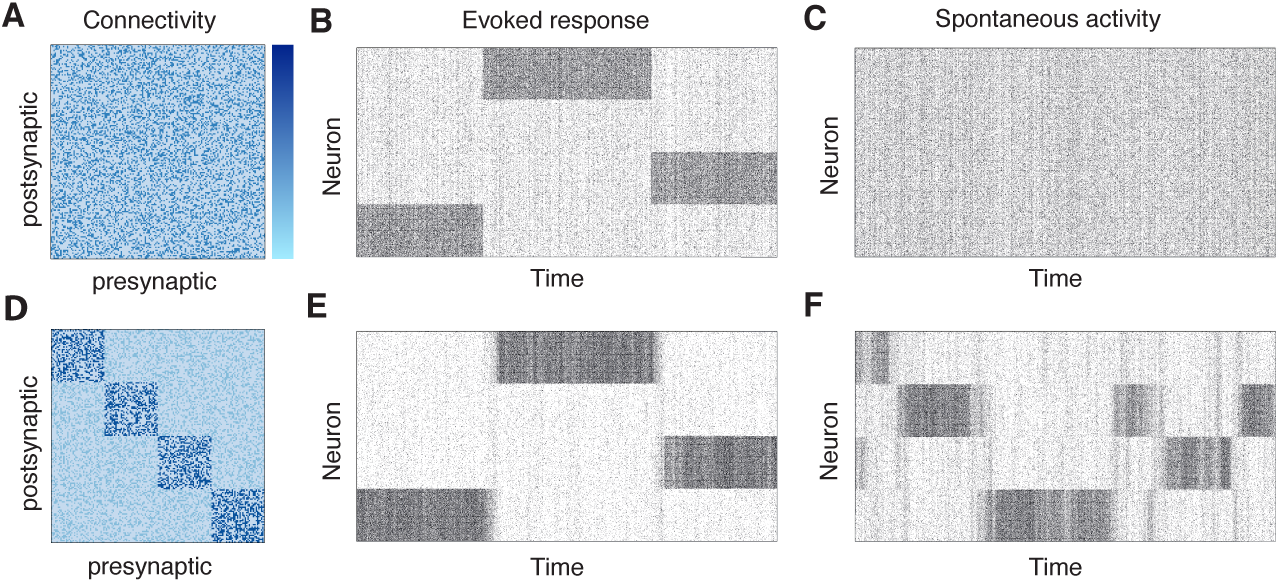
**A.** Excitatory connectivity matrix of an unstructured recurrent network of excitatory and inhibitory spiking neurons [52^•^, 57]. **B.** Spike rasters of the evoked response in the network by driving three different subsets of excitatory neurons with stronger external input compared to the other neurons, as indicated by the elevated firing rates. **C.** Activity in the network in response to uniform external input to all excitatory neurons. **D.** Excitatory connectivity matrix of a structured recurrent network of excitatory and inhibitory spiking neurons. Neurons are more strongly connected within a cluster, which could be imprinted through plasticity mechanisms in simulated networks [54^•^, 55^•^, 56]. **E.** Spike rasters of the evoked response as in B. **F.** In response to uniform external input to all excitatory neurons, the network spontaneously activates subsets of neurons with stronger connectivity [55^•^, 56]. These could be interpreted as attractors of the network dynamics, giving rise to spontaneous retrieval of evoked activity patterns. This behavior is absent in the unstructured network (C).

Innovative theoretical analysis has also derived the conditions for the spontaneous emergence of different types of assemblies through STDP at recurrent synapses, in the absence of feedforward patterned inputs [58]. By changing the shape of the plasticity rule and the biophysical properties of synaptic transmission, the authors demonstrated the emergence of self-connected assemblies vs. synfire chains [58]. Interestingly, the same structures emerged upon training with random vs. temporal sequences of inputs (respectively) in models with feedforward and recurrent plasticity under voltage STDP [44].

The development of functional recurrent circuitry in these models often relies on an interplay of Hebbian and homeostatic forms of plasticity. Classical Hebbian-style plasticity rules alone induce a positive feedback instability whereby neurons that are frequently co-active will increase their connectivity and future co-activity. To combat this problem and bring circuit function to a normal operating regime, the above models implement diverse homeostatic mechanisms based on experimental observations [59]: (1) normalization of synaptic weights, (2) metaplasticity where the amplitude and sign of Hebbian synaptic change is modulated ((1) and (2) reviewed in [60^••^]) (3) plasticity at inhibitory synapses [54^•^, 55^•^, 56] and (4) shifts in intrinsic excitability [61^•^, 62], or a combination of them [63, 64^•^]. A key insight from these models has been that experimental forms of homeostatic plasticity are too slow to stabilize Hebbian plasticity; stability in the models requires faster forms of homeostatic plasticity that have yet to be identified experimentally [60^••^, 65].

Taken together, these studies highlight the importance of theory and models to understand how functional connectivity in recurrent networks emerges from Hebbian and homeostatic plasticity giving rise to stable dynamics and computations. A future challenge would be to interpret these findings in the context of specific biophysical mechanisms that might implement them (e.g. [66]), and to relate them to the detailed map formation models discussed earlier [67^•^]. Moreover, it would be interesting to examine the emergence of functional organization under realistic developmental patterns of activity, which as discussed earlier are sluggish and might utilize different plasticity rules than those that rely on precise spike timing [68].

## Conclusion

Theoretical and computational approaches have contributed in powerful ways to our understanding of how neuronal circuits develop to establish precise connectivity and tuned single neuron responses, and to give rise to adult computations. Retinotopic map formation represents perhaps the most successful example of models of development (apart from orientation maps): starting from phenomenological models, theorists have proposed more comprehensive models which can explain larger data sets and make interesting predictions. However, this represents only one aspect of neural development. Going forward, we should use this example to build modeling frameworks which capture the diversity of mechanisms unique to this period, their timescales and spatial scales of operation and their coordinated action to generate adult computations.

In addition to the detailed analysis of spontaneous and sensory-evoked activity in developing circuits *in vitro*, we still need to understand the generation and function of this activity in the intact animal. With the recent spur of in vivo recordings [26^•^, 50^•^, 24, 25^•^], theoretical neuroscience can contribute to the quantitative analysis of longitudinal recordings of single neuron and network activity in novel ways. This analysis can provide us with necessary assumptions and constraints for new models of how this activity is generated, how it changes over development, and what its role is in sculpting developing networks.

Analyzing this activity can also help us infer the appropriate developmental plasticity rules from the potentially different correlational structure in the juvenile and the adult [38, 39, 69]. This will enable us to link theoretical descriptions of plasticity at the level of neuron pairs to network connectivity refinements, explaining the emergence of functional units such as synfire chains, assemblies and memory attractors [54^•^, 55^•^, 58]. The observation that the same network structures emerge either intrinsically through the properties of the plasticity rule, or externally through the nature of the input patterns, suggests that these issues should be examined experimentally under specific developmental scenarios where the derived model structures are observed.

While it seems natural that models should explore novel hypotheses and make predictions to direct future experiments, we also point out another important role. Existing models should be tested on paradigms and data different from those on which the models were initially based. This has the value of testing the generality and utility of modes and avoids overfitting. Theory and models hold the potential to uncover common underlying principles (or differences) in the development of different circuits, for instance sensory and motor [70^•^]. In some cases, the same solution might emerge for different problems, but often different solutions might be beneficial to ensure robustness and variability of responses.

With the accumulation of experimental data, theory and models need to play a larger role in understanding the development of neural circuits with its diversity of interacting instructive signals guiding self-organization. We have proposed that the new focus should be on the developmental emergence of single cell properties and population activity patterns, and the behaviorally relevant computations these might reflect. As many developmental processes are carefully orchestrated, theoretical and modeling approaches are necessary to tease apart the relative importance and role of each process.

## Acknowledgements

This work was supported by funding from the Max Planck Society. JG holds a Career Award at the Scientific Interface from the Burroughs Wellcome Fund. The authors thank Matthias Kaschube, Tatjana Tchumatchenko and Stephen Eglen for comments on the manuscript.

## References

[1] Koulakov AA and Tsigankov DN. A stochastic model for retinocollicular map development. BMC Neurosci, 5:30, 2004.

[2] Eglen SJ and Gjorgjieva J. Self-organization in the developing nervous system: theoretical models. HFSP Journal, 3:176–185, 2009.

[3] van Ooyen A. Using theoretical models to analyse neural development. Nat Rev Neurosci, 12:311–326, 2011.

[4] Goodhill GJ. Contributions of theoretical modeling to the understanding of neural map development. Neuron, 56:301–311, 2007.

[5] Cang J and Feldheim DA. Developmental mechanisms of topographic map formation and alignment. Annu Rev Neurosci, 36:51–77, 2013.

[6] Goodhill GJ. Can molecular gradients wire the brain? Trends Neurosci, 39:202–211, 2016.

[7] Reingruber J and Holcman D. Computational and mathematical methods for morphogenetic gradient analysis, boundary formation and axon targeting. Semin Cell Dev Biol, 35:189–202, 2014. • This review discusses models and algorithms of the interplay between morphogenetic gradients and patterned activity aimed towards bridging the gap between molecular instructions and large-scale, system-level organization.

[8] Thompson A, Gribizis A, Chen C, and Crair MC. Activity-dependent development of visual receptive fields. Curr Opin Neurobiol, 42:136–143, 2017. •• A detailed review discussing aspects of receptive field development and plasticity in the lateral geniculate nucleus, primary visual cortex and superior colliculus instructed by spontaneous and visually driven activity.

[9] Huberman AD, Feller MB, and Chapman B. Mechanisms underlying development of visual maps and receptive fields. Annu Rev Neurosci, 31:479–509, 2008.

[10] Grimbert F and Cang J. New model of retinocollicular mapping predicts the mechanisms of axonal competition and explains the role of reverse molecular signaling during development. J Neurosci, 32:9755–9768, 2012.

[11] Godfrey KB, Eglen SJ, and Swindale NV. A multi-component model of the developing retinocollicular pathway incorporating axonal and synaptic growth. PLoS Comput Biol, 5:e1000600, 2009.

[12] Godfrey KB and Swindale NV. Modeling development in retinal afferents: retinotopy, segregation and EphrinA/EphA mutants. PLoS One, 9:e104670, 2014. •• This work extends the model from [11]. Besides being able to generate morphologically realistic axonal growth patterns observed experimentally during map formation, this extended model also includes the emergence of several receptive field properties, for example, the segregation of retinal axons based on their origin eye (left or right) and their stimulus polarity preference (ON or OFF).

[13] Willshaw DJ, Sterratt DC, and Teriadikis A. Analysis of local and global topographic order in mouse retinocollicular maps. J Neurosci, 34:1791–1805, 2014.

[14] Owens MT, Feldheim DA, Stryker MP, and Triplett JW. Stochastic interactions between neural activity and molecular cues in the formation of topographic maps. Neuron, 87:1261–1273, 2015.

[15] Hjorth JJJ, Sterratt DC, Cutts CS, Willshaw DJ, and Eglen SJ. Quantitative assessment of computational models for retinotopic map formation. Dev Neurobiol, 75:641–666, 2014. •• The authors provide a simulation framework to assess existing models of map formation (focusing on retinocollicular maps) in a quantitative and unbiased manner and compare them against experimental data. The authors also provide the simulation software.

[16] Tikidji-Hamburyan RA, El-Ghazawi RA, and Triplett JW. Novel models of visual topographic map alignment in the superior colliculus. PLoS Comput Biol, 12:e1005315, 2016. •• Proposes two alternative models for how the topographic maps between the retina and the SC on one hand, and V1 and the SC on another, align in the SC. The first is a ‘correlation model’ which assumes that the SC is strongly driven by the retina only, and the alignment with V1 happens though correlation-based firing mechanisms. The second is an ‘integration model’ which assumes that V1 inputs also drive SC firing during alignment. While both models reproduce *in vivo* data, the authors make novel predictions that would distinguish between them.

[17] Moody WJ and Bosma MM. Ion channel development, spontaneous activity and activity dependent development in nerve and muscle cells. Physiol Rev, 85:883–941, 2005.

[18] Kirkby LA, Sack GS, Firl A, and Feller MB. A role for correlated spontaneous activity in the assembly of neural circuits. Neuron, 80:1129–1144, 2013.

[19] Ackman JB and Crair MC. Role of emergent neural activity in visual map development. Curr Opin Neurobiol, 24:166–175, 2014.

[20] Blankenship AG and Feller MB. Mechanisms underlying spontaneous patterned activity in developing neural circuits. Nat Rev Neurosci, 11:18–29, 2010.

[21] Lansdell B, Ford K, and Kutz JN. A reaction-diffusion model of cholinergic retinal waves. PLoS Comput Biol, 10:e1003953, 2014.

[22] Gjorgjieva J and Eglen SJ. Modeling developmental patterns of spontaneous activity. Curr Opin Neurobiol, 21:679–684, 2011.

[23] Colonnese MT and Khazipov R. ’Slow activity transients’ in infant rat visual cortex: a spreading synchronous oscillation patterned by retinal waves. J Neurosci, 30:4325–4337, 2010.

[24] Siegel F, Heimel AJ, Peters J, and Lohmann C. Peripheral and central inputs shape network dynamics in the developing visual cortex in vivo. Curr Biol, 22:253–258, 2012.

[25] Ackman JB, Burbridge TJ, and Crair MC. Retinal waves coordinate patterned activity throughout the developing visual system. Nature, 490:219–225, 2012. • This tour-de-force *in vivo* imaging of spontaneous activity in the mouse developing visual system shows that spontaneous wave of retinal activity propagate throughout the entire visual system before eye opening. Retinal waves propagate in the superior colliculus, lateral geniculate nucleus and primary visual cortex. In contrast, secondary visual areas are only modulated by retina wave activity. Therefore, the study postulates the role of this activity for driving connectivity refinements along the entire visual pathway.

[26] Shen J and Colonnese MT. Development of activity in mouse visual cortex. J Neurosci, 36:12259–12275, 2016. • This experimental study gives a comprehensive timeline of spontaneous activity and evoked responses in developing mouse visual cortex. By the time of eye-opening spontaneous activity is indistinguishable from mature activity. Multiple aspects of evoked responses are also studied.

[27] Moreno-Juan V, Filipchuk A, Antón-Bolaños N, Mezzera C, Gezelius H, Andrés B, Rodríguez-Malmierca L, Susín R, Schaad O, Iwasato T, Schüle R, Rutlin M, Nelson S, Ducret S, Valdeolmillos M, Rijli FM, and López-Bendito G. Prenatal thalamic waves regulate cortical area size prior to sensory processing. Nat Commun, 8:14172, 2017.

[28] Mease RA, Famulare M, Gjorgjieva J, Moody WJ, and Fairhall A. Emergence of adaptive computation by single neurons in the developing cortex. The Journal of Neuroscience, 33:12154–12170, 2013.

[29] Gjorgjieva J, Mease RA, Moody WJ, and Fairhall AL. Intrinsic neuronal properties switch the mode of information transmission in networks. PLoS Comput Biol, 10:e1003962, 2014. •• Using simulation and analysis, this study shows how the early development of biophysical properties of cortical cells impacts the functional properties at the level of networks. As single neurons become able to scale the gain of their responses to the amplitude of the stimuli they encounter, a property termed ‘gain scaling’ (see [28]), networks of these neurons transition from being able to generate large-scale propagating wave events to being efficient encoders of fast information. The gain scaling ability is linked to the membrane conductances in the neuron model, which are developmentally regulated.

[30] Baltz T, Herzog A, and Voigt T. Slow oscillating population activity in developing cortical networks: models and experimental results. J Neurophysiol, 106:1500–1514, 2011.

[31] Barnett HM, Gjorgjieva J, Weir K, Comfort C, Fairhall AL, and Moody WJ. Relationship between individual neuron and network spontaneous activity in developing mouse cortex. J Neurophysiol, 112:3033–3045, 2014. • Early cerebral cortex shows propagating waves of spontaneous activity. Ventrolateral piriform cortex is uniquely able to initiate such waves. The study examines these waves in slice experiments, and shows through modeling that the experimental results can be generated in a network with recurrent synaptic connectivity and local gap junctions, and an increasing gradient of intrinsic excitability at the level of single neurons from piriform to neocortex.

[32] Ben-Ari Y. Excitatory actions of GABA during development: the nature of the nurture. Nat Rev Neurosci, 3:728–739, 2002.

[33] Ben-Ari Y, Gaiarsa J-L, Tyzio R, and Khazipov R. GABA: A pioneer transmitter that excites immature neurons and generates primitive oscillations. Physiol Rev, 87:1215–1284, 2007.

[34] Kanold PO and Luhmann HJ. The subplate and early cortical circuits. Annu Rev Neurosci, 33:23–48, 2010.

[35] Viswanathan S, Sheikh A, and Looger LL Kanold PO. Molecularly defined subplate neurons project both to thalamocortical recipient layers and thalamus. Cereb Cortex, in press, 2016.

[36] Murata Y and Colonnese MT. An excitatory cortical feedback loop gates retinal wave transmission in rodent thalamus. eLife, 5:e18816, 2016. •• The authors show the existence of a transient circuit in the feedback from cortex to thalamus. Early corticothalamic feedback acts excitatory on thalamus through gap junctions amplifying thalamic activity driven by spontaneous retinal waves. This transient feedback ensures high-fidelity transmission of sensory signals when connections are immature and switches to the adult structure of corticothalamic inhibition just prior to eye-opening to prevent large scale epileptic activity in the adult.

[37] Kanold PO and Shatz CJ. Subplate neurons regulate maturation of cortical inhibition and outcome of ocular dominance plasticity. Neuron, 51:627–638, 2006.

[38] Butts DA, Kanold PO, and Shatz CJ. A burst-based “Hebbian” learning rule at retino-geniculate synapses links retinal waves to activity-dependent refinement. PLoS Biol, 5:e61, 2007.

[39] Gjorgjieva J, Toyoizumi T, and Eglen SJ. Burst-Time-Dependent Plasticity robustly guides ON/OFF segregation in the Lateral Geniculate Nucleus. PLoS Comput Biol, 5:e1000618, 2009.

[40] Dilger EK, Krahe TE, Morhardt DR, Seabrook TA, Shin H-S, and Guido W. Absence of plateau potentials in dLGN cells leads to a breakdown in retinogeniculate refinement. J Neurosci, 35:3652–3662, 2015.

[41] Luhmann HJ, Sinning A, Yang J-W, Reyes-Puerta V, Stüttgen MC, Kirischuk S, and Kilb W. Spontaneous neuronal activity in developing neocortical networks: From single cells to large-scale interactions. Front Neural Circuits, 10:40:doi:10.3389/fncir.2016.00040, 2016.

[42] Bi GQ and Poo MM. Synaptic modifications in cultured hippocampal neurons: dependence on spike timing, synaptic strength, and postsynaptic cell type. J Neurosci, 18:10464–10472, 1998.

[43] Pfister J-P and Gerstner W. Triplets of spikes in a model of spike timing-dependent plasticity. J Neurosci, 26:9673–9682, 2006.

[44] Clopath C, Büsing L, Vasilaki E, and Gerstner W. Connectivity reflects coding: A model of voltage based STDP with homeostasis. Nat Neurosci, 13:344–352, 2010.

[45] Gjorgjieva J, Clopath C, Audet J, and Pfister J-P. A triplet spike-timing-dependent plasticity model generalizes the Bienenstock-Cooper-Munro rule to higher order spatiotemporal correlations. Proc Natl Acad Sci USA, 108:19383–19388, 2011.

[46] van Hooser SD, Escobar GM, Maffei A, and Miller P. Emerging feed-forward inhibition allows the robust formation of direction selectivity in the developing ferret visual cortex. J Neurophysiol, 111:2355–2373, 2014.

[47] Brito CSN and Gerstner W. Nonlinear Hebbian learning as a unifying principle in receptive field formation. PLoS Comput Biol, 12:e1005070, 2016. • Various approaches for imprinting receptive fields, both normative and biophysical, implement a common generic principle: non-linear Hebbian learning, which is defined as the effective weight change curve arising from the combination of the functional form of the learning window and the neurons input-output curve.

[48] Olshausen BA and Field DJ. Emergence of simple-cell receptive field properties by learning a sparse code for natural images. Nature, 381:607–609, 1996.

[49] Bell AJ and Sejnowski TJ. The “Independent Components” of natural scenes are edge filters. Vision Res, 37:3327–3338, 1997.

[50] Smith GB, Sederberg A, Elyada YM, van Hooser SD, Kaschube M, and Fitzpatrick D. The development of cortical circuits for motion discrimination. Nat Neurosci, 18:252–261, 2015. • Beyond selectivity, discrimination of stimuli depends on the variability of responses. The authors show that activity in ferret cortex at eye opening expresses high levels of activity that spread wave-like, high variability and strong noise correlations. After three weeks, activity is sparser, and both variability and noise correlations are decreased considerably.

[51] Ko H, Cossell L, Baragli C, Antolik J, Clopath C, Hofer SB, and Mrsic-Flogel TD. The emergence of functional microcircuits in visual cortex. Nature, 496:96–100, 2013. •• Using experiment, the authors show that neurons in mouse visual cortex are tuned at eye-opening but that connections in the network are not specifically stronger between neurons with similar response properties than between neurons with dissimilar responses, as it is the case in adult cortex. Through modeling they show how such specificity of connections can develop after eye-opening through synaptic plasticity, and how the initial tuning can arise in early stages of development.

[52] Sadeh S, Clopath C, and Rotter S. Emergence of functional specificity in balanced networks with synaptic plasticity. PLoS Comput Biol, 11:e1004307, 2015. • The authors show that STDP at excitatory and inhibitory synapses in random networks with dominant inhibition gives rise to tuning-specific connection strength between neurons, further sharpening the orientation selectivity and extending the contrast invariance apparent in such networks.

[53] Rochefort NL, Garaschuk O, Milos R-I, Narushima M, Marandi N, Pichler B, Kovalchuk Y, and Konnerth A. Sparsification of neuronal activity in the visual cortex at eye-opening. Proc Natl Acad Sci USA, 106:15049–15054, 2009.

[54] Miconi T, McKinstry JL, and Edelman GM. Spontaneous emergence of fast attractor dynamics in a model of developing primary visual cortex. Nat Commun, 7:13208, 2015. • The specific achievement of this study is the simultaneous emergence of feedforward receptive fields and recurrent assembly structure through plasticity in networks which are driven by inputs emulating natural image statistics.

[55] Litwin-Kumar A and Doiron B. Formation and maintenance of neuronal assemblies through synaptic plasticity. Nat Commun, 5:5319, 2014. • The study investigates the emergence of preferentially connected groups of neurons in recurrent networks through training, and how stable this imprinted structure is without further training and when training the network with a new set of stimuli.

[56] Zenke F, Everton JA, and Gerstner W. Diverse synaptic plasticity mechanisms orchestrated to form and retrieve memories in spiking neural networks. Nat Commun, 6:6922, 2015.

[57] Amit DJ and Brunel N. Model of spontaneous activity and local structured activity during delay periods in the Cerebral Cortex. Cereb Cortex, 7:237–252, 1997.

[58] Ravid Tannenbaum N and Burak Y. Shaping neural circuits by high-order synaptic interactions. PLoS Comput Biol, 12:e1005056, 2016.

[59] Turrigiano GG and Nelson SB. Homeostatic plasticity in the developing nervous system. Nat Rev Neurosci, 5:97–107, 2004.

[60] Zenke F and Gerstner W. Hebbian plasticity requires compensatory processes on multiple timescales. Philos Trans R Soc B Biol Sci, 372:20160259, 2017. •• This review discusses theoretical results on the timescales on which homeostatic plasticity is required to act in order to stabilize the dynamics of neural systems and the discrepancy between these theoretically desirable and the experimentally measured timescales on which known homeostatic mechanisms act.

[61] Harnack D, Pelko M, Chaillet A, Chitour Y, and van Rossum MCW. Stability of neuronal networks with homeostatic regulation. PLoS Comput Biol, 11:e1004357, 2015. • A systematic analysis of the necessary conditions for stable homeostatic control of intrinsic excitability in both single neurons and recurrent networks. Homeostatic control interacts with the strength of recurrent couplings and the timescale of activity, both of which constrain the timescale on which homeostasis can act without destabilizing the system.

[62] Cannon J and Miller P. Synaptic and intrinsic homeostasis cooperate to optimize single neuron response properties and tune integrator circuits. J Neurophysiol, 116:2004–2022, 2016.

[63] Lazar A, Pipa G, and Triesch J. SORN: a self-organizing recurrent neural network. Front Comput Neurosci, 3:23, 2009.

[64] Zheng P, Dimitrakakis C, and Triesch J. Network self-organization explains the statistics and dynamics of synaptic connection strength in cortex. PLoS Comput Biol, 9:e1002848, 2013. • This particular study on SORNs shows that self-organization in networks equipped with excitatory spike timing dependent plasticity, structural plasticity, and three forms of homeostatic plasticity, synaptic scaling, intrinsic plasticity of firing thresholds and inhibitory spike timing dependent plasticity, gives rise to dynamics and distribution of synaptic efficacies resembling experimental findings.

[65] Toyoizumi T, Kaneko M, Stryker MP, and Miller KD. Modeling the dynamic interaction of Hebbian and homeostatic plasticity. Neuron, 84:497–510, 2014.

[66] Graupner M and Brunel N. Calcium-based plasticity model explains sensitivity of synaptic changes to spike pattern, rate, and dendritic location. Proc Natl Acad Sci USA, 109:3991–3996, 2012.

[67] Sweeney Y and Clopath C. Emergent spatial synaptic structure from diffusive plasticity. Eur J Neurosci, doi:10.1111/ejn.13279, 2016. • The proposed model simultaneously established map-like structures and the emergence of strongly interconnected groups in response to correlated external drive. This was accomplished by incorporating nitric oxide (a diffusive neurotransmitter that can alter the excitability of other nearby neurons even in the absence of synaptic coupling) into the Bienenstock-Cooper-Munro rule.

[68] Graupner M, Wallisch P, and Ostojic S. Natural firing patterns imply low sensitivity of synaptic plasticity to spike timing compared with firing rate. J Neurosci, 36:11238–11258, 2016.

[69] Lim S, McKee JL, Woloszyn L, Amit Y, Freedman DJ, Sheinberg DL, and Brunel N. Inferring learning rules from distributions of firing rates in cortical neurons. Nat Neurosci, 18:1804–1810, 2015.

[70] Gjorgjieva J, Evers JF, and Eglen SJ. Homeostatic activity-dependent tuning of recurrent networks for robust propagation of activity. J Neurosci, 36:3722–3734, 2016. • This modeling study shows how networks of excitatory and inhibitory populations assemble through plasticity and spontaneous activity to produce sequentially propagating activity. It is shown that homeostatic plasticity maintaining postsynaptic firing rates in a stable range (and not Hebbian plasticity) robustly tunes network connectivity to produce propagating rhythmic activity, as it is seen e.g. in *Drosophila* larvae motor network during crawling.

